# FSCN1 is critical for HNSCC

**DOI:** 10.1101/2023.06.20.545739

**Authors:** Xin Wei

## Abstract

**Background:** The study of molecular markers for diagnosis and prognosis is of great clinical significance for HNSCC patients. In this study, we proposed that FSCN1 has a potential indication for prognosis and is essential for the migration of HNSCC.

**Methods:** We analyzed the expression and survival association of FSCN1 in HNSCC using TCGA data. We compared the expression of FSCN1 in tumors from primary and metastasis HNSCC patients using QPCR, western blotting, and immunochemistry staining. We determined the migration velocity of multiple HNSCC cell lines using a chemotaxis migration assay. We analyzed the correlation between FSCN1 expression and HNSCC cell migration. We also test the effect of FSCN1 knockdown and overexpression on HNSCC cell migration.

**Results:** FSCN1 was overexpressed in HNSCC than pair normal tissues and metastasis HNSCC than primary HNSCC. FSCN1 expression was associated with significantly poorer overall survival of HNSCC patients. FSCN1 was potentially associated with immune cell infiltration and migration-associated genes. FSCN1 level was correlated with the migration in HNSCC cell lines. Knockdown of FSCN1 reduced the migration and the overexpression of FSCN1 promoted the migration of HNSCC cell lines.

**Conclusion:** FSCN1 is a potential prognostic marker and a critical biomolecule for the migration of HNSCC

## 1. Introduction

Head and neck squamous cell carcinoma (HNSCC) is the sixth most common cancer in the world. Each year, over 300,000 new HNSCC cases occur in the world and result in about 3 hundred thousand deaths globally [1]. HNSCC, including laryngeal cancer, oral cancer, nasopharyngeal cancer, and hypopharyngeal cancer, has many primary sites and pathological types and more than 90% of head and neck tumors are squamous cell carcinoma [2]. Although surgical treatment, radiotherapy, and chemotherapy have been applied for the treatment of HNSCC, the prognosis of HNSCC currently is unfavourable [3, 4]. One of the most challenging issues in HNSCC treatment is the diagnosis and prognosis of HNSCC [4]. If the condition is not diagnosed and treated in a timely manner, it can lead to lymph node metastasis and result in a higher mortality rate for patients [5]. Therefore, the study of molecular markers for diagnosis and prognostic is of great clinical significance for HNSCC patients. The identification of metastasis regulators for HNSCC is conducive to the prevention of HNSCC metastasis and the development of HNSCC therapy.

FSCN1, Fascin Actin-Bundling Protein 1, is an actin-bundling protein that is suggested to affect cell migration [6]. Fascin proteins can organize F-actin into parallel bundles therefore they are the key proteins for the formation of actin-based cellular protrusions [7-9]. In addition, FSCN1 also functions in the organization of actin filament bundles and the formation of microspikes, membrane ruffles, and stress fibres [10]. Studies have shown that FSCN1 is critical for the formation of a diverse set of cell protrusions, such as filopodia, and cell motility and migration [7-9]. FSCN1 was also supposed to mediate the reorganization of the actin cytoskeleton and axon growth cone collapse in response to the nerve growth factor [10]. Therefore, we suggested that FSCN1 might be one of the key biomolecules for cancer cell migration, motility, adhesion, and cellular interactions.

In cancer, the FSCN1 expression level was proposed to associate with clinically pathogenic phenotypes in multiple cancers type [6]. It has been reported that the overexpression or silencing of FSCN1 affected both the invasion and migration of multiple types of cancer cell lines, such as squamous cell carcinoma [11-13] and adenocarcinoma [14-16]. A study revealed that the expression of FSCN1 in hepatocellular carcinoma was associated with the loss of matrix metalloproteinases [17]. However, few studies have been reporting the effect of FSCN1 on HNSCC migrations.

In this study, we tested FSCN1 as a prognostic biomarker for clinical HNSCC patients. We proposed that FSCN1 is essential for the migration of HNSCC cells. Thus, this study provided a novel molecular biomarker for the prognosis and migration regulator of HNSCC.

## 2. Methods

### 2.1. The acquisition of the transcriptome data

The mRNA sequencing data of HNSCC samples were accessed from The Cancer Genome Atlas (TCGA) in January 2020, following the required guidelines and policies.

### 2.2. Bioinformatic analysis

ggplot2 (v3.3.2) was used to conduct an analysis of the genes and survival with R foundation for statistical computing (2020) version 4.0.3. The correlation of HSCN1 expression and migration states in HNSCC single-cell datasets GSE103322 were accessed and analyzed using the CancerSEA.

### 2.3. The acquisition of clinical HNSCC samples

HNSCC tissue samples were collected from 29 patients including 14 patients with primary HNSCC and 15 patients with HNSCC metastasis. All donors were older than 18 years old and have been informed and consented to the use of the samples.

### 2.4. Cell culture

A-253 and SCC-25 (SC-25) were purchased from ATCC (Washington, USA). BICR-31 (ICR-31) was purchased from Ximbio (London, UK). YD-38 were the oral cancer cell lines derived from oral squamous cell carcinoma tissue from patients [18]. BICR-22 and BICR-6 were purchased from the ECACC (Salisbury, UK). CAL-33 and BICR-56 were purchased from the DSMZ (Braunschweig, Germany). All cells were cultured in DMEM medium with the supplement of 2mM Glutamine and 10% FBS in a cell culture incubator of 5% CO2 and 37°C.

### 2.5. Cell transfection

knockdown and overexpression of FSCN1 in cells were conducted by transfection of FSCN1 shRNA or FSCN1 expression plasmids into cells. The predesigned FSCN1 shRNA (FSCN1 Human shRNA Lentiviral Particle Locus ID 6624) plasmid and Control (Lenti particles Scrambled shRNA), Human FSCN1 expression plasmid (NM_003088), and packaging plasmids Lenti-vpak packaging kit with transfection reagent (TR30037) were purchased from OriGene (Rockville, MD, USA) and the experiments were conducted following the instruction of the kit.

### 2.6. QPCR

The mRNA levels of FSCN1 in samples were determined using a QPCR assay[19]. The RNeasy Mini kit (Qiagen, Germantown, MD, USA) was used to extract RNA from samples. The PrimeScript RT Reagent kit with gDNA Eraser (Takara Bio, Japan) was used to retro transcription RNA into DNA. The PowerUp™ SYBR™ Green Master Mix (Thermo, Beverly, MA, USA) was used to conduct the QPCR with the Applied Biosystems StepOnePlus instrument (Thermo, Beverly, MA, USA). The FSCN1 expression was normalized with GAPDH expression using the 2-ΔΔCT method. The sequences of the primers were as follows: FSCN1 forward: 5’-GGCAAGTTTGTGACCTCCAAGAA-3’; FSCN1 reverse: 5’-AGCCGATGAAGCCATGCTC-3’; GAPDH forward 5’ -ACAACTTTGGTATCGTGGAAGG-3’; GAPDH reverse: 5’-GCCATCACGCCACAGTTTC-3’.

### 2.7. Western blotting

The protein expression of FSCN1 was analyzed using a western blotting assay. The Pierce protein lysing buffer (Rockford, IL, USA) with Roche protease inhibitors were used to extract proteins in samples. Then 10–12% sodium dodecyl sulfate-polyacrylamide gels were used to separate the proteins. The proteins were then transferred to nitrocellulose membranes for immune reaction. The membrane was blocked in the blocking buffer (5% milk) for 1 h, then incubated with the primary antibodies overnight at 4 °C following the incubation of the secondary antibodies at room temperature for 1 h. ECL reagent was used to visualize the secondary antibodies and analysis of the protein level. The FSCN1 protein expression was normalized with GAPDH protein expression. Antibodies used for western were (sc-21743) Anti-Fascin 1 Antibody (55K-2), Anti-GAPDH antibody [6C5]-Loading Control (ab8245), and Rabbit Anti-Mouse IgG H&L (HRP) (ab6728).

### 2.8. Immunochemistry staining

FSCN1 staining was done by immunochemistry. Paraffin-embedded tissue samples were deparaffinized in xylene and rehydrated through graded ethanols. These samples were then submerged into the citric acid buffer for heat-induced antigenic retrieval. All the samples were then blocked with 10% BSA to reduce the background. Then, samples were incubated with FSCN1 primary antibodies at 4°C overnight and the color of the sample was developed using the DAKO ChemMate Envision Kit HRP (Dako-Cytomation, Carpinteria, CA, USA) followed by counterstaining with hematoxylin, dehydration, clearing and mounting.

### 2.9. Chemotaxis assay

The m-Slide chemotaxis system (ibidi, Germany) was used to conduced the real-time cell migration assay for individual cells [20, 21]. Cells were cultured on the central channel of the chemotaxis slide at 10% confluency. The images of migrating cells were recorded for 10 hours using a time-lapse Micro-Imager and the tracks of the individual cell were analyzed. A computer with the Ibidi software was used to analyze the velocity of cell migration.

### 2.10. Statistics of bench works

All the results were presented in X ± SD in the figures. A t-test or ANOVA with Dunnett’s post hoc tests was used to analyze the significance of the differences (p < 0.05). The GraphPad Prism (version 8) was used to plot the charts and calculate statistics.

## 3. Results

### 3.1. FSCN1 has a potential indication for the prognosis of HNSCC

In this study, we explored the prognostic value of FSCN1 for clinical HNSCC. We first compared the FSCN1 level in TCGA HNSCC transcriptome and paired normal tissues. Results showed that FSCN1 was significantly overexpressed in HNSCC compared to paired normal tissues (Fig.1A). This indicated that FSCN1 was potentially useful for the clinical diagnosis of HNSCC. We also analyzed the potential association between FSCN1 expression and the overall survival of HNSCC patients. Results revealed that FSCN1 was significantly associated with survival with a p-value of 0.002 when we did a log-rank analysis comparing 0-50% low and 50-100% high FSCN1 patients (Fig.1B). Univariate Cox regression further demonstrated that FSCN1 was a risk factor for HNSCC patients with a hazard ratio of 1.54. Multivariate Cox regression analysis revealed the FSCN1 was independent of age and pTNM stages, two factors that affected overall survival in the univariate Cox regression result (Fig.1C). According to the multivariate Cox regression analysis, a nomogram was constructed (Fig.1D). The nomogram aimed for the prediction of 1-, 3-, 5-year survival for HNSCC patients. We then calculated calibration curves for the nomogram. The grey diagonal line on the calibration curve of the overall survival nomogram model in the discovery group represents the ideal nomogram, while the red, yellow, and blue lines represent the 1-year, 3-year, and 5-year of the observed nomogram, respectively. The red, yellow, and blue lines were all close to the grey line, indicating that the predicted overall survival was consistent with the observed survival data. This demonstrates that the nomogram is reliable and can be used to accurately predict patient outcomes (Fig.E). The nomogram demonstrated the possibility of clinical application of FSCN1 in HNSCC prognosis.

**Figure 1.**
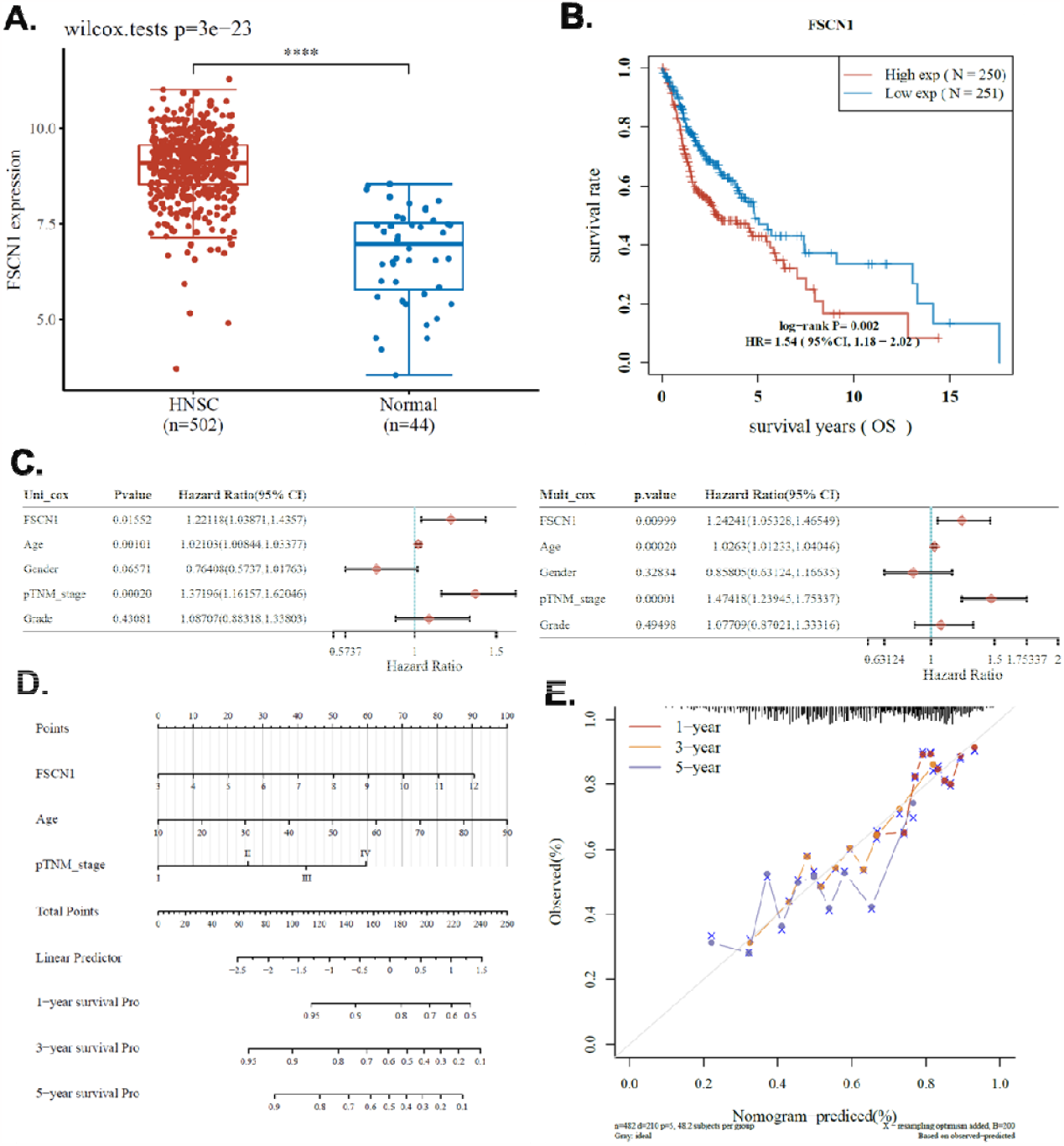
The prognostic power of FSCN1. A. The expression level of the FSCN1 gene in HNSCC and pair normal tissue from TCGA HNSCC dataset. B. The survival of patients with high and low FSCN1 expressed. C. Univariate and multivariate Cox regression analysis of FSCN1 expression and common clinical factors. D. Nomogram constructed according to the multivariate regression analysis. E. Calibration curve of the nomogram.

### 3.2. Associated genes enrichment analysis

To obtain information on the potential mechanisms involved in the effect of the FSCN1 gene on HNSCC, we identified differentially expressed genes (DEGs) between FSCN1 high (75-100%) and low (0-25%) groups. We set the cutoff fold change value of 2 and the p-value of 0.05. Results showed that 393 (up) and 322 (down) genes were identified as DEGs positively and negatively associated with FSCN1 in HNSCC respectively (Fig.2A-B). These genes were further enriched in GO terminologies and KEGG pathways. Results of KEGG enrichment showed that genes positively associated with FSCN1 were enriched in “PI3K−Akt signaling pathway” and “Focal adhesion”, while genes negatively associated with FSCN1 were enriched in “Cytokine−cytokine receptor interaction”. In terms of GO enrichment, genes positively associated with FSCN1 were enriched in “extracellular structure organization”, “extracellular matrix organization”, “epidermis development”, and “skin development”, while genes negatively associated with FSCN1 were enriched in “lymphocyte differentiation”, “T cell activation”, and “regulation of cell−cell adhesion” (Fig.2C).

**Figure 2.**
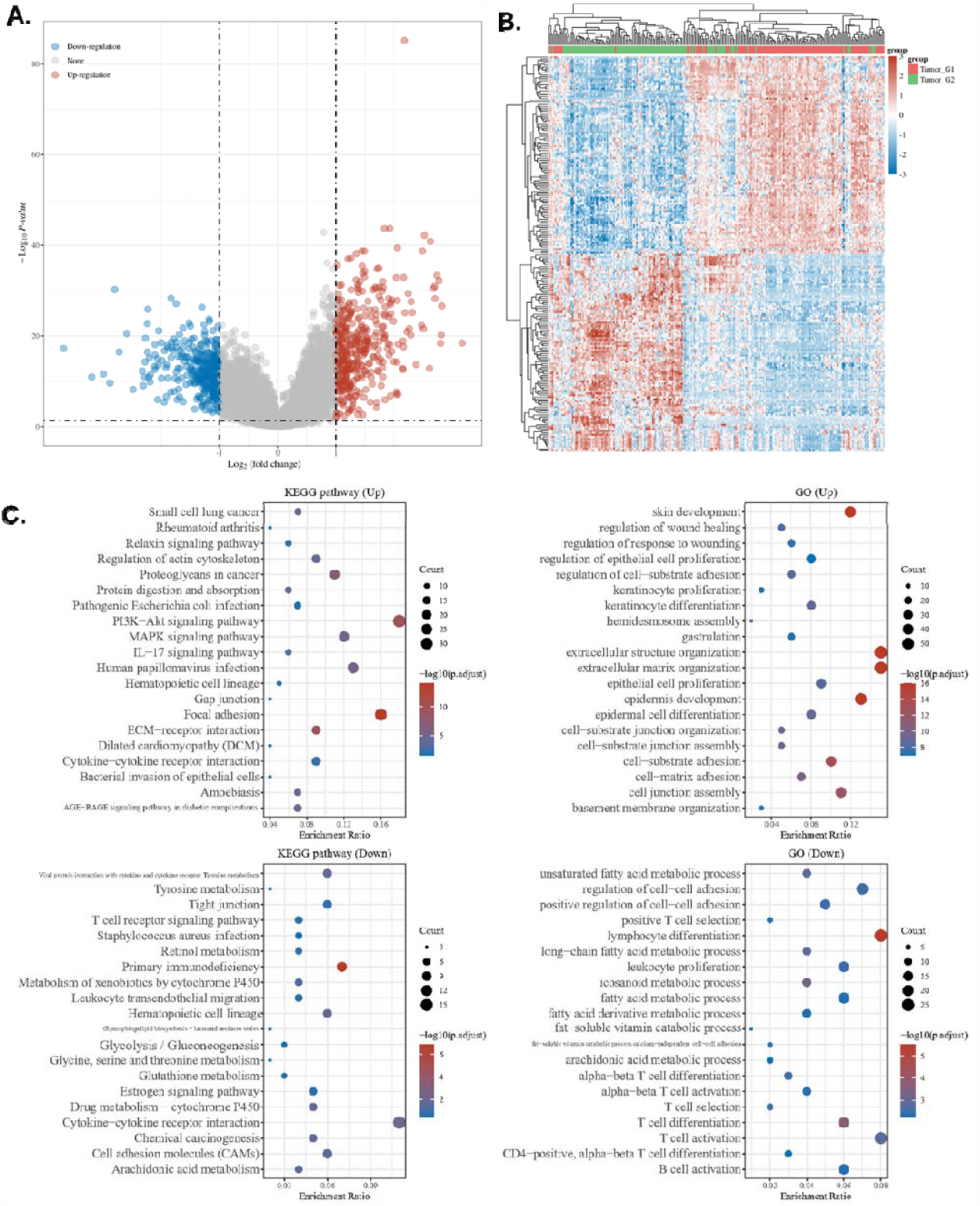
Potential association of FSCN1 and HNSCC migration. A-C. Associated genes enrichment analysis. A-B. Differentially expressed gene analysis in HNSCC FSCN1. FSCN1 high (75-100%) and low (0-25%) samples were compared. A. Volcano plots were constructed using fold-change >2 and P<0.01. The red spots in the plot represent the gene positively associated with FSCN1 and the blue spots represent the gene negatively associated with FSCN1. B. Heat map and hierarchical clustering analysis of FSCN1 associated genes. C. The GO and the KEGG signaling pathways enrichment analysis of FSCN1 associated genes.

### 3.3. Potential association of FSCN1 and immune cell infiltration

We conducted the Spearman correlation analysis of FSCN1 expression and immune cell infiltration of HNSCC using the EPIC algorithms. Results showed that FSCN1 expression was strongly and positively correlated with T cell CD4+ and negatively correlated with B cell and T cell CD8+ (Fig.3A). These results suggested that in high FSCN1 expressing HNSCC, the tumor had a higher level of T cell CD4+ and a lower level of B cell and T cell CD8+. Although our results were not able to validate the functional role of FSCN1 in the regulation of tumor immune microenvironment in HNSCC, the expression of HNSCC is a biological sign of the changes in the immune response.

**Figure 3.**
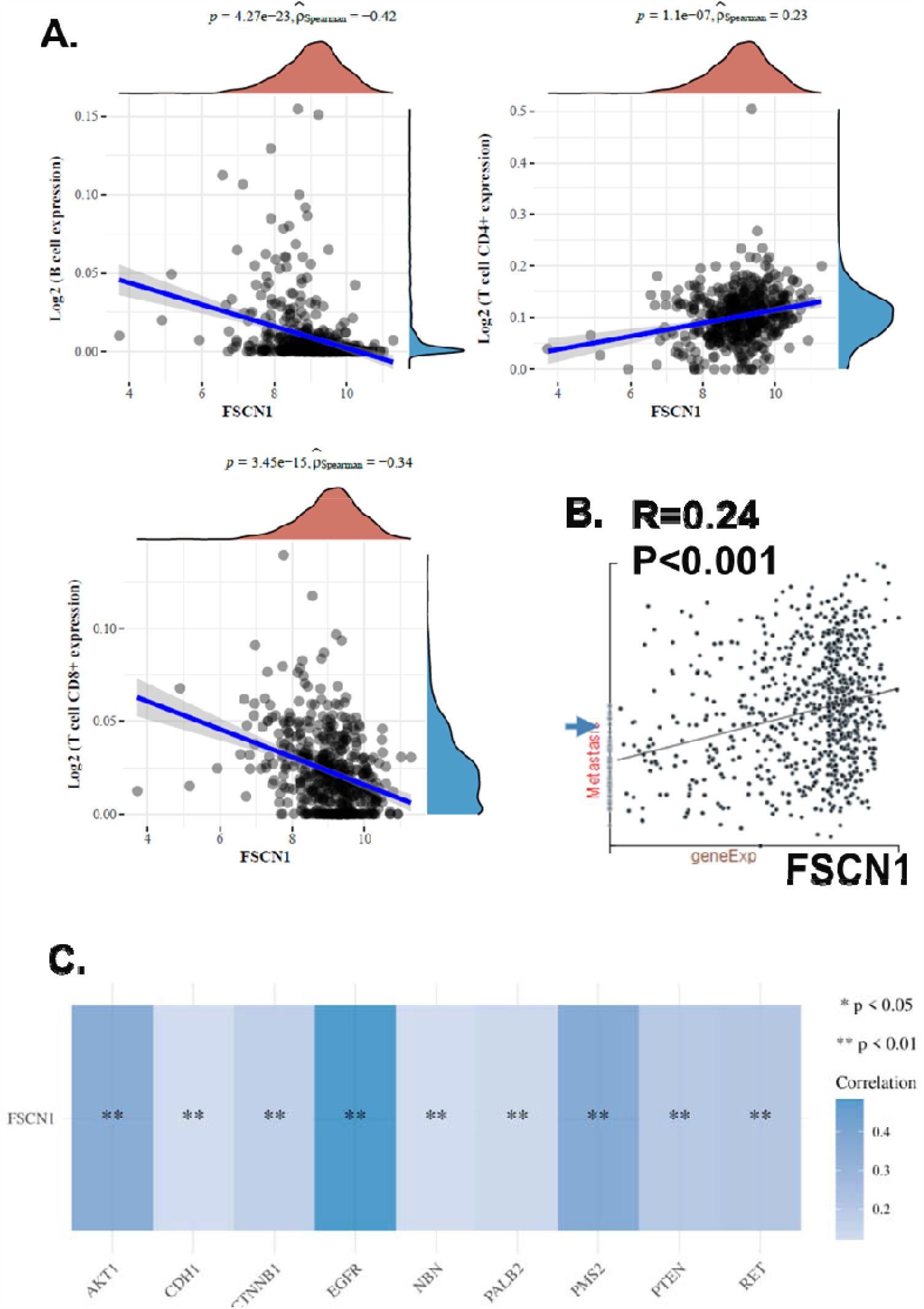
Potential association of FSCN1 and immune/migration of HNSCC. A. The correlations between FSCN1 expression and immune cell infiltration levels in HNSCC. B. The correlations between FSCN1 expression and migration state in HNSCC single-cell datasets. The GSE103322 data (n=2105) were accessed and analyzed using the CancerSEA. C. The correlations between FSCN1 expression and common migration-associated genes using TCGA HNSCC dataset.

### 3.4. Potential association of FSCN1 and HNSCC migration

To explore the potential role of FSCN1 in HNSCC migration, we investigated a single-cell data set of HNSCC using the CancerSEA. We analyzed the correlation of FSCN1 expression and HNSCC migration state scores of 2150 single HNSCC cells. Results showed that FSCN1 was positively correlated with HNSCC migration with a coefficient of 0.24 (Fig.3B). On the other hand, we analyzed the correlation of FSCN1 and several common HNSCC migration-associated genes, including EGFR, PMS2, AKT1, RET, PTEN, CTNNB1, PALB2, NBN, and CDH1. Results showed that EGFR, PMS2, AKT1, RET, PTEN, CTNNB1, PALB2, NBN, and CDH1 were positively associated with FSCN1 with coefficients of 0.48, 0.37, 0.35, 0.21, 0.20, 0.17, 0.14, 0.12, and 0.12 respectively (Fig.3C). Therefore, we suggested FSCN1 might be associated with HNSCC migration.

### 3.5. FSCN1 expression difference between metastasis and primary HNSCC

To further investigate the role of FSCN1 in HNSCC migration, we collected cancer tissue samples from 14 patients with primary HNSCC and 15 patients with HNSCC metastasis. We conducted QPCR, western blotting, and immunochemistry staining to compare FSCN1 expression levels in metastasis and primary HNSCC samples. Results showed that metastasis HNSCC samples had a higher level of FSCN1 at both mRNA and protein levels (Fig.4A-C). Immunochemistry staining showed that metastasis HNSCC was strongly stained while the primary HNSCC staining was weak. The staining results further validated that FSCN1 is highly expressed in metastasis HNSCC compared with the primary HNSCC. We suggested that the metastasis association of FSCN1 might account for the potential indication for prognosis.

**Figure 4.**
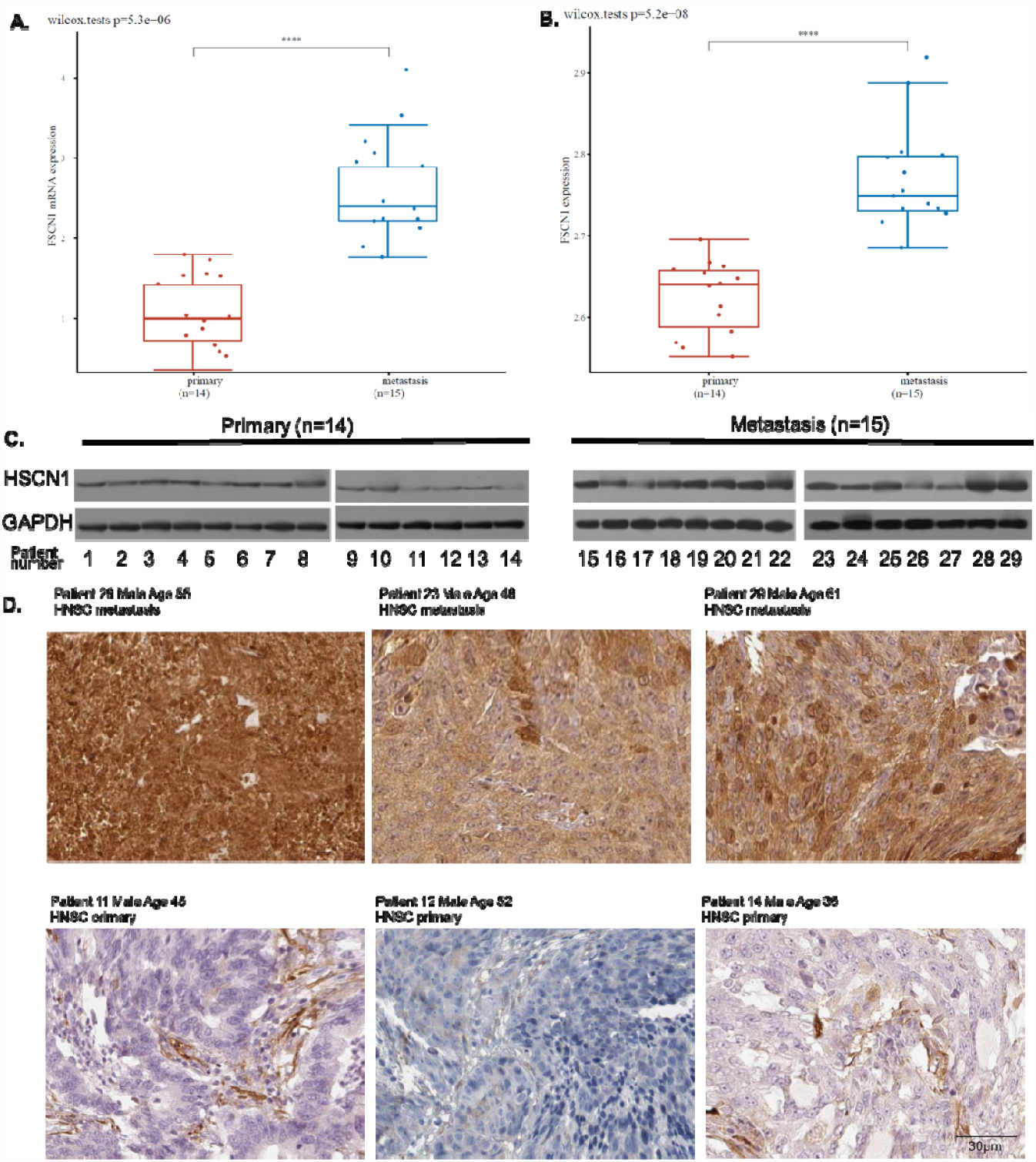
FSCN1 expression difference between metastasis and primary HNSCC. A. The mRNA expression of FSCN1 in metastasis and primary HNSCC samples (QPCR). B. The protein expression of FSCN1 in metastasis and primary HNSCC samples (western blotting). C. Representative image of the western blotting of tumor tissues from primary(n=14) or metastasis(n=15) samples. D. Representative image of protein staining of FSCN1 in metastasis and primary HNSCC samples.

### 3.6. Correlation of FSCN1 expression and HNSCC cell line migration

In the subsequent study, we designed experiments using HNSCC cell models. We first screened proper cell lines for the study. We plotted the FSCN1 expression in 33 HNSCC cell lines in the CCLE database. We found that all these 33 cell lines expressed FSCN1 (Fig.5A). The protein of FSCN1 is mainly localized to the Cytosol of cells. Then we validated the FSCN1 levels in eight of these cell lines using QPCR and western blotting in our lab (Fig.5B&C&D). These eight cell lines included A-253, CAL-33, ICR-31, SC-25, YD-38, BICR-22, BICR-6, and BICR-56. Western blotting showed that A-253 and CAL-33 had the highest expression of FSCN1 while BICR-22, BICR-6, and BICR-56 had the lowest expression of FSCN1. The QPCR results showed a similar trend as the western blotting. Next, we conducted in vitro chemotaxis assay to observe the migration of individual HNSCC cells. The computer recorded the track of the cell every 10 seconds and calculated the velocity of cell migration. Results showed that CAL-33 and ICR-31 were the fastest migrating cell lines while BICR-56 was the slowest migrating cell line. We further plotted the correlation between the protein expression of FSCN1 and the velocity of eight HNSCC cell lines. Results showed that the protein expression of FSCN1 is significantly correlated with the velocity of eight HNSCC cell lines(Fig.6). We suggested that the impact of FSCN1 on cancer cells might account for the potential indication for prognosis.

**Figure 5.**
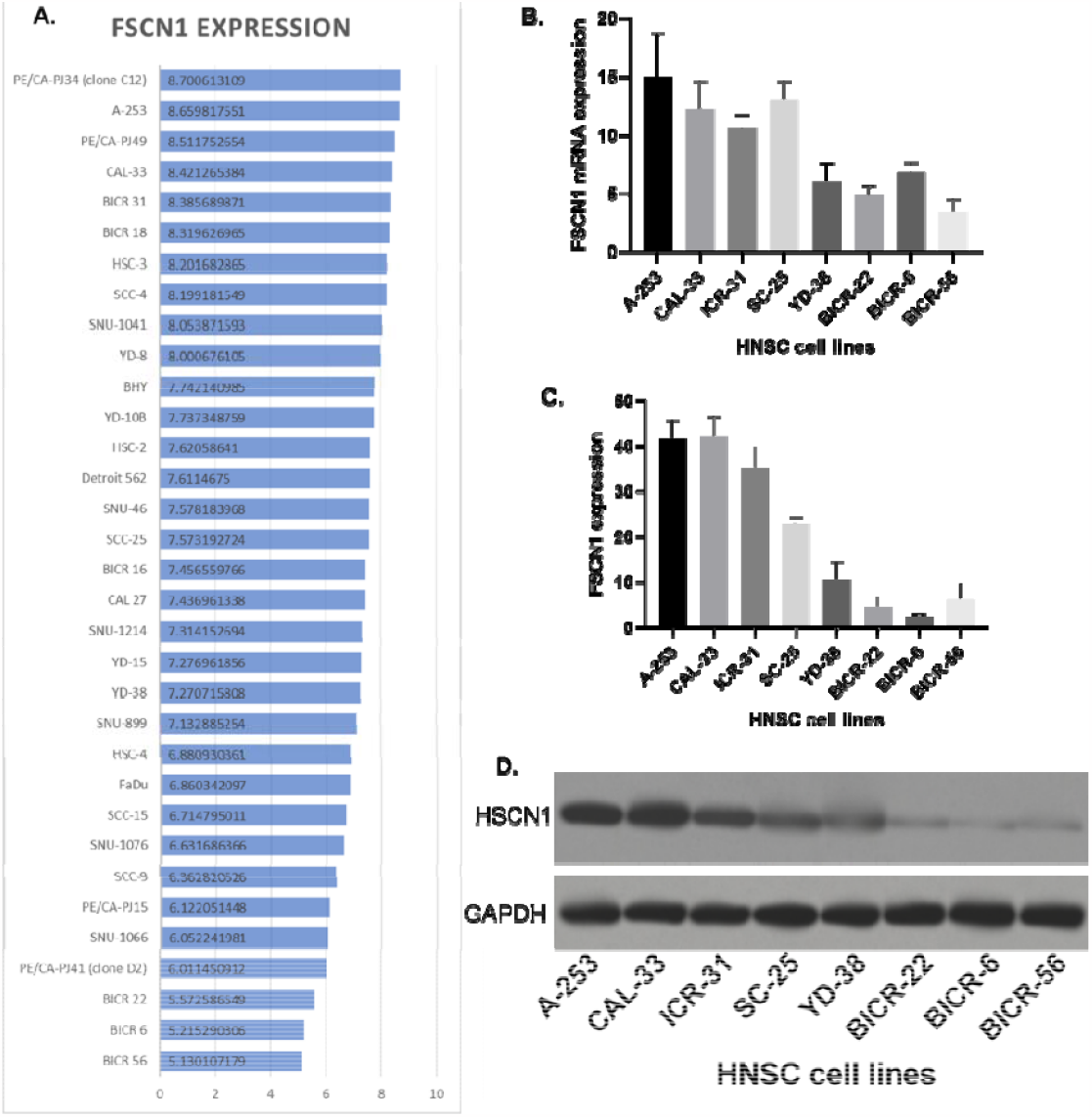
The expression of FSCN1 in HNSCC cell lines. A. The expression level of FSCN1 in HNSCC cell lines from the CCLE dataset. B. The mRNA expression of FSCN1 in eight HNSCC cell line samples (QPCR). C. The protein expression of FSCN1 in eight HNSCC cell lines samples (western blotting). D. Image of the western blotting.

**Figure 6.**
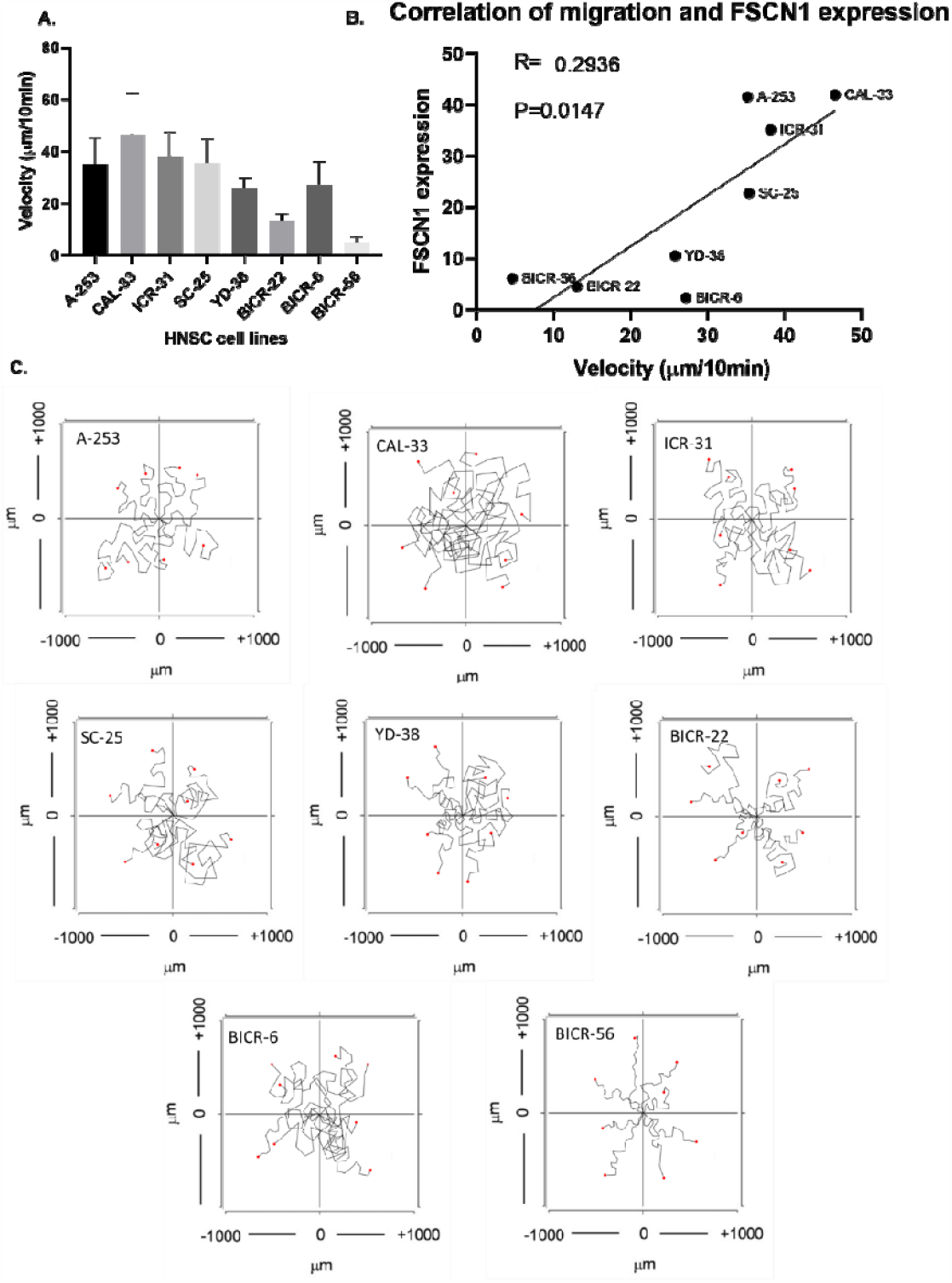
Correlation of FSCN1 expression and HNSCC cell line migration. A. The velocity of eight HNSCC cell lines. B. Correlation of the protein expression of FSCN1 and the velocity of eight HNSCC cell lines. C. Images of the single-cell migration track.

### 3.7. The functional role of FSCN1 in HNSCC migration

To investigate whether FSCN1 was functional in HNSCC migration regulations, we overexpressed FSCN1 in BICR-6 and BICR-56 where FSCN1 is expressed lowly. Results showed that 20 nM of FSCN1 expressing plasmid highly improved the levels of FSCN1 in BICR-6 and BICR-56 cells (Fig.7A-B). The overexpression of FSCN1 using 20 nM of FSCN1 overexpression plasmid significantly increased the velocity of the migration in both BICR-6 and BICR-56 cell lines (Fig.7E-G). In addition, we also knocked down FSCN1 expression in two of the most FSCN1-expressing cell lines CAL-33 and A253. Results showed that 20 nM of FSCN1 shRNA plasmid almost completely block the expression of FSCN1 in CAL-33 and decreased FSCN1 expression in A253 cells by over 90%. The chemotaxis assay results showed that the knockdown of FSCN1 significantly reduced the velocity of the migrations in both A253 and CAL-33 cell lines(Fig.8).

**Figure 7.**
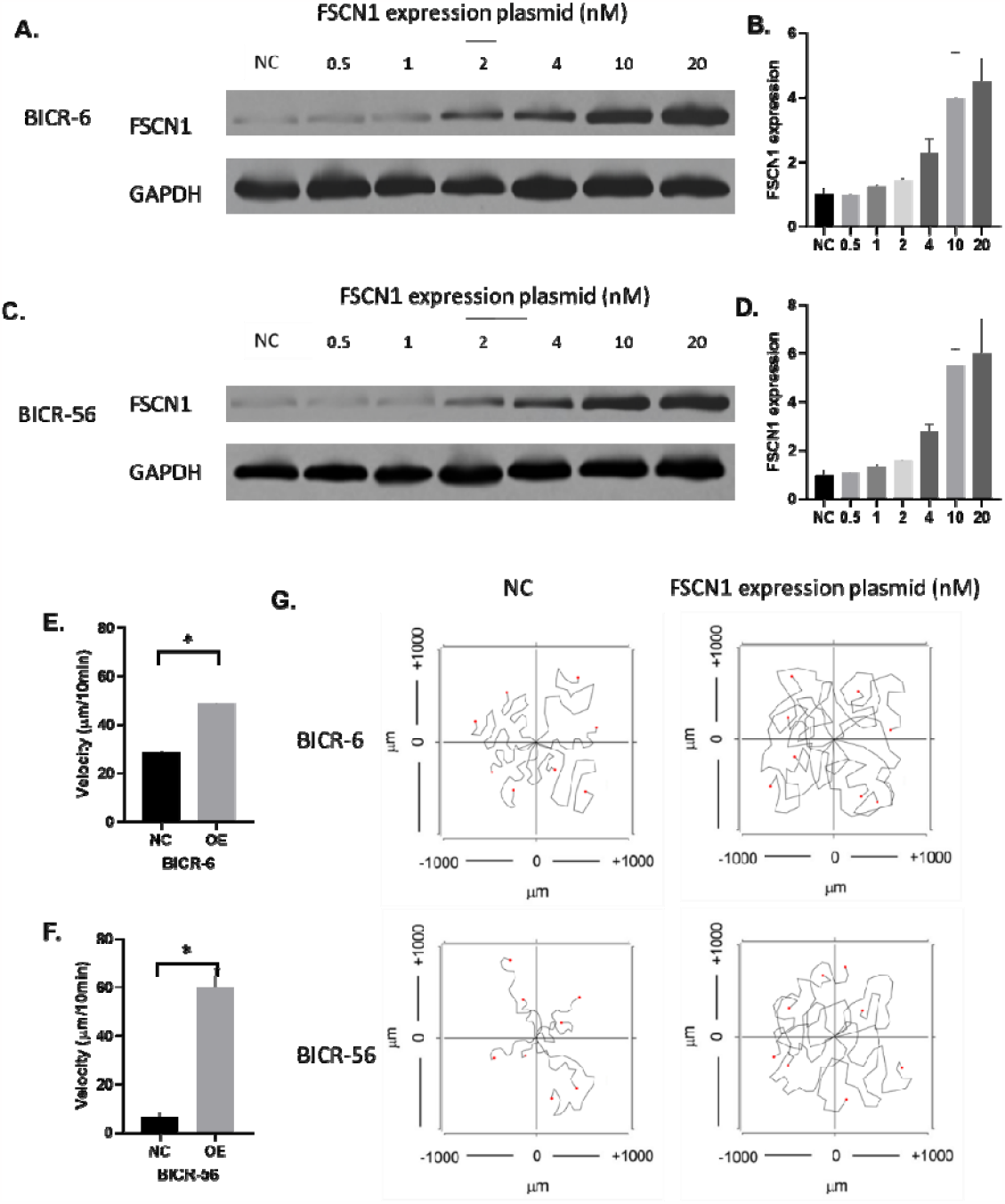
The overexpression of FSCN1 in BICR-6 and BICR-56. A-B. The protein expression of FSCN1 in BICR-6 with different levels of FSCN1 overexpression. C-D. The protein expression of FSCN1 in BICR-56 samples with different levels of FSCN1 overexpression. E-F. The velocity of BICR-6 and BICR-56 with and without FSCN1 overexpression. G. Images of the single-cell migration track of BICR-6 and BICR-56.

**Figure 8.**
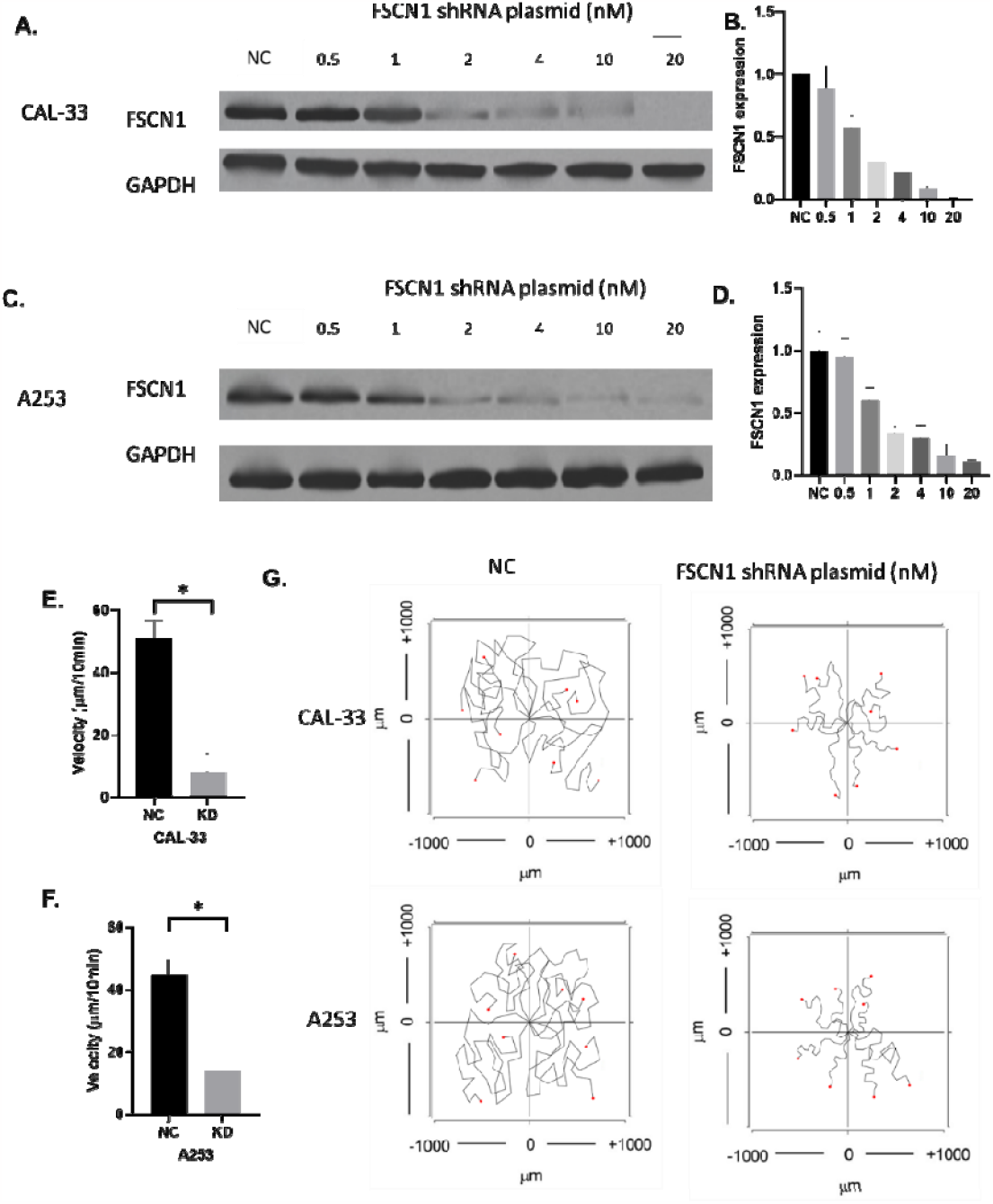
The knockdown of FSCN1 in CAL-33 and A253. A-B. The protein expression of FSCN1 in CAL-33 with different levels of FSCN1 knockdown. C-D. The protein expression of FSCN1 in A253 samples with different levels of FSCN1 knockdown. E-F. The velocity of CAL-33 and A253 with and without FSCN1 knockdown. G. Images of the single-cell migration track of CAL-33 and A253.

## 4. Discussion

In clinical treatment, HNSCC patients with recurrence and/or metastasis have an extremely low survival rate: the median overall survival rate is only 1 year [22], thus, the metastasis and migration of HNSCC cells have been one of the most concerns for HNSCC treatment. In this study, we demonstrated that FSCN1 was a reliable biomarker for HNSCC diagnosis and prognosis. Bioinformatic studies have been wildly used to access the survival associations of genes[4, 23-33]. Our data suggested that the overall survival rate of HNSCC patients was significantly associated with the expression of FSCN1 in HNSCC tissues. The prognostic potential of FSCN1 has been reported in other types of cancers. In colon cancer, FSCN1 has been found to be positively correlated with survival[34]. The FSCN1 gene was also suggested as an important predictor of early-stage breast cancer [35]. In addition, FSCN1 expression was associated with significantly poorer overall survival of epithelial ovarian cancer patients [36]. In renal cell carcinoma, FSCN1 was overexpressed in tumor tissues compared to non-tumor tissues. It is also associated with poor overall survival and recurrence-free survival in renal cell carcinoma, [37]. In all of these analyses, high expression of FSCN1 was associated with a worse prognosis and most of the studies revealed that FSCN1 expression was independent of age, tumor size, and clinical TNM stage, which were consistent with our results. As for the subtype of HNSCC, a study focused on HNSCC in the tongue found that FSCN1 is an effective biomarker of poor prognosis and a potential therapeutic target in human tongue HNSCC[38]. In our study, we study all types of HNSCC as a whole and further demonstrated that the FSCN1 was not only useful for tongue HNSCC but also for other HNSCC. All these bioinformatic studies demonstrated that FSCN1 might be valuable for clinical cancer prognosis. Nevertheless, we suggested that it is not certain whether this molecule has clinical or therapeutic relevance and can be applied for clinical HNSCC treatment. Our study did not test the therapeutic relevance and the clinical part of the study only provided an observation, not necessarily proving that it can be a biomarker and potentially be better than current biomarkers. Therefore, we should be cautious when drawing any conclusions.

A previous study reported FSCN1 as a potential therapeutic target of lung squamous cell carcinoma and suggested that FSCN1 impact immune and inflammation in the tumor microenvironment[39]. As another type of squamous cell carcinoma, HNSCC might have a similar impact on the immune and inflammation in the tumor microenvironment as lung cancer. In this study, we found that FSCN1 was potentially associated with B cells and T cell infiltration levels. Our results did not validate the functional role of FSCN1 in the regulation of tumor immune microenvironment in HNSCC. However, its expression in HNSCC could still be a biological sign of the changes in the immune response. The changes in immune cell infiltration levels can contribute to the metastasis of the HNSCC. The tumor extracellular matrix (ECM) of HNSCC is significantly different from that of normal tissue[40]. The ECM components upregulated in HNSCC can impact several cancer hallmarks, such as sustaining proliferative signaling, promoting angiogenesis, facilitating invasion and metastasis, modulating growth suppressor activity, and suppressing antitumor immunity. Furthermore, the tumor ECM has been implicated in treatment resistance, making it a potential therapeutic target. Malignant epithelial cells and stromal cells in HNSCC interact through signaling, which leads to the increased production of components of the ECM. These ECM components provide a substrate for carcinoma cell migration, modulate the cytokine environment, and aid in immune evasion. Therefore, we suggested that, as an actin-binding protein mediating the formation of actin-based cellular protrusions, FSCN1 might be involved in the ECM pathways thereby affecting tumor microenvironment and immune cell infiltration levels. In clinical treatment, many different therapies might be applied to inhibit cancer cells [41]. In cancer treatment, many clinical factors might affect cancer metastasis by either regulating cancer cells or the immune system such as the use of anaesthetics or traditional medicines [42-48]. The potential regulation of FSCN1 on immune cells might contribute to the effects of these factors on cancers. Yet, further studies are required to identify the direct impact of FSCN1 on immune cells.

Moreover, we not only analyzed data from the TCGA but also collected clinical samples and validated the expression. Commonly used QPCR and western blotting assay were conducted in many previous studies [49, 50], which provided reliable results. We also used immunochemistry staining [51] to visualize the expression of FSCN1 in HNSCC tissues. One of the most significant findings of this study was that FSCN1 was found to be the essential molecule for HNSCC cell migration. It was not surprising that FSCN1 potentially affected cancer cell migration because FSCN1 has long been defined as a fascin actin-bundling protein that is responsible for the migration or relocation of cells [7-9]. Animal models have been widely used in research previously [52-54]. For the FSCN1 study, an in vivo study using the orthotopic xenografts mice model indicated that FSCN1 could promote cancer cell invasion and metastasis [37]. Another study revealed that FSCN1 could affect gastric cancer migration and invasion [55]. However, so far, few studies have focused on the role of FSCN1 in HNSCC migration.

We notice that a previous study reported that FSCN1 is an effective marker of poor prognosis and a potential therapeutic target in human tongue squamous cell carcinoma[38], yet, the previous study focused on tongue squamous cell carcinoma, while our study expanded the application of FSCN1 to general HNSCN. In addition, we have used multiple cancer cell lines to support the role of FSCN1 in cancer migration, which has never been done before. The correlation of HSCN1 expression and velocity of migration of the eight HNSCC cell lines we tested strongly suggested that FSCN1 was a key molecule for HNSCC cell migration. We validated the promoting effect of FSCN1 on HNSCC cell migration by overexpressing FSCN1 in two of the low FSCN1 cell lines with low migration velocity. We further confirmed the functional effect of FSCN1 by knocking down FSCN1 in two of the high FSCN1 cell lines with high migration velocity. These results were consistent with the previous conclusion that FSCN1 regulated cancer cell migration and metastasis. Furthermore, most of the previous studies investigated cancer cell migration using wound-healing or transwell assay, which determined the migration of a group of cancer cells as a whole. Our study was the first to investigate the role of FSCN1 by tracking individual cell migration. Nevertheless, although we obtain the velocity of individual cell migration, we were not able to determine the expression of FSCN1 in the cell we recorded.

This study aimed to validate the role of FSCN1 in HNSCC cell migration, but the mechanisms underlying these effects were not clear. The enrichment analysis revealed several potential mechanisms, such as the PI3K−Akt signaling pathway, which required further validation with experimental evidence. In addition, as the FSCN1 might play a role in intercellular communications, many cancer-related ion channels, such as VGSC[56-58], TRP[59, 60], and TPCs[61, 62], might also contribute to the effects of FSCN1 on cancer cells and need further investigation in the future.

## 5. Conclusion

FSCN1 is a potential prognostic marker and a critical biomolecule for the migration of HNSCC.

## 6. Declarations

### Ethics approval and consent to participate

The study has been approved by the Ethics Committee of Hainan General Hospital and consents to participate.

### Consent for publication

Not applicable.

### Availability of data and materials

Raw data and materials are available from the corresponding author by reasonable request.

### Authors’ contributions

CONCEPTION: Yuliang Zhang and Anyan Zhou

INTERPRETATION OR ANALYSIS OF DATA: Yuliang Zhang and Anyan Zhou

PREPARATION OF THE MANUSCRIPT: Yuliang Zhang and Anyan Zhou

REVISION FOR IMPORTANT INTELLECTUAL CONTENT: Jiabin Nian, Shuzhou Liu, and Xin Wei

SUPERVISION: Xin Wei

## Acknowledgments

None.

